# A harmonized meta-knowledgebase of clinical interpretations of cancer genomic variants

**DOI:** 10.1101/366856

**Authors:** Alex H Wagner, Brian Walsh, Georgia Mayfield, David Tamborero, Dmitriy Sonkin, Kilannin Krysiak, Jordi Deu Pons, Ryan P Duren, Jianjiong Gao, Julie McMurry, Sara Patterson, Catherine Del Vecchio Fitz, Ozman U Sezerman, Jeremy L Warner, Damian T Rieke, Tero Aittokallio, Ethan Cerami, Deborah Ritter, Lynn M Schriml, Robert R Freimuth, Melissa Haendel, Gordana Raca, Subha Madhavan, Michael Baudis, Jacques S Beckmann, Rodrigo Dienstmann, Debyani Chakravarty, Xuan Shirley Li, Susan Mockus, Olivier Elemento, Nikolaus Schultz, Nuria Lopez-Bigas, Mark Lawler, Jeremy Goecks, Malachi Griffith, Obi L Griffith, Adam A Margolin, Variant Interpretation for Cancer Consortium

## Abstract

Precision oncology relies on the accurate discovery and interpretation of genomic variants to enable individualized diagnosis, prognosis, and therapy selection. We found that knowledgebases containing clinical interpretations of somatic cancer variants are highly disparate in interpretation content, structure, and supporting primary literature, impeding consensus when evaluating variants and their relevance in a clinical setting. With the cooperation of experts of the Global Alliance for Genomics and Health (GA4GH) and six prominent cancer variant knowledgebases, we developed a framework for aggregating and harmonizing variant interpretations to produce a meta-knowledgebase of 12,856 aggregate interpretations covering 3,437 unique variants in 415 genes, 357 diseases, and 791 drugs. We demonstrated large gains in overlap between resources across variants, diseases, and drugs as a result of this harmonization. We subsequently demonstrated improved matching between a patient cohort and harmonized interpretations of potential clinical significance, observing an increase from an average of 33% per individual knowledgebase to 56% in aggregate. Our analyses illuminate the need for open, interoperable sharing of variant interpretation data. We also provide an open and freely available web interface (search.cancervariants.org) for exploring the harmonized interpretations from these six knowledgebases.

---

The promise of precision oncology–in which a cancer patient’s treatment is informed by the mutational profile of their tumor–requires concise, standardized, and searchable clinical interpretations of the detected variants. These structured interpretations of biomarker-disease associations can be diagnostic (determinant of a disease type or subtype), prognostic (indicator of patient outcome), therapeutic (predictive of favorable or adverse response to therapy), or predisposing (germline variants that increase risk of developing cancer). Isolated institutional efforts have contributed to the curation of the biomedical literature to collect and formalize these interpretations into knowledgebases^1–12^. These isolated efforts have resulted in disparate models for representing this knowledge, and the rapid generation and evolution of knowledge compounds this heterogeneity. While the vast scale of the curation process drives the need for many individual efforts, the heterogeneity we face when exchanging biomarker-disease associations represents a critical challenge that must be addressed^13^. Consequently, stakeholders interested in the effects of genomic variants of a cancer on potential therapeutic interventions are faced with the following tradeoff: 1) referencing and understanding multiple representations and interpretations of variants across knowledgebases; or 2) potentially omitting clinically significant interpretations that are not universally captured across knowledgebases. Manual aggregation of information across knowledgebases to interpret each patient’s variant profile is an unsustainable approach that does not scale in a precision medicine setting. Moreover, the lack of an integrated resource has precluded the ability to assess the current state of precision treatment options based on the aggregated knowledge across major cancer precision medicine programs. Published reports^14–17^ have relied on individual, often highly discordant knowledgebases. Interoperability and automated aggregation is therefore required to make a comprehensive approach to cancer precision medicine tractable and to compare interpretations across knowledgebases in order to establish consensus.

The current diversity and number of “knowledge silos” and the associated difficulties of coordinating these disparate knowledgebases has led to an international effort to maximize genomic data sharing.^18, 19^ The Global Alliance for Genomics and Health (GA4GH) has emerged as an international cooperative project to accelerate the development of approaches for responsible, voluntary, and secure sharing of genomic and clinical data.^20, 21^ The Variant Interpretation for Cancer Consortium (VICC; cancervariants.org) is a *Driver Project* of GA4GH, established to co-develop standards for genomic data sharing (ga4gh.org/howwework/driver-projects.html). Specifically, the VICC is a consortium of clinical variant interpretation experts addressing the challenges of representing and sharing curated interpretations across the cancer research community.

In this study, we leveraged the VICC member expertise to aggregate cancer variant interpretations from six distinguished constituent knowledgebases: Cancer Genome Interpreter (CGI), Clinical Interpretations of Variants in Cancers (CIViC), Jackson Labs Clinical Knowledgebase (JAX-CKB), MolecularMatch, OncoKB, and the Precision Medicine Knowledgebase (PMKB) (**Table S1**).^1, 5, 9–11, 22^ The institutions leading each constituent knowledgebase agreed upon a core set of principles (http://cancervariants.org/principles/) stipulating that the contents of each knowledgebase would contain a minimum set of data elements describing an interpretation and be freely shared with the research community.

This cooperative effort enabled us to develop a framework for structuring and harmonizing clinical interpretations across these knowledgebases. Specifically, we defined key elements of variant interpretations (genes, variants, diseases, drugs, and evidence), developed strategies for harmonization (through linking these elements to established and unambiguous references), and implemented this framework to consolidate interpretations into a single, harmonized meta-knowledgebase (freely available at search.cancervariants.org).

## RESULTS

### A strategy for aggregating and structuring interpretation knowledge

An initial survey of the constituent knowledgebases of the VICC (**Table S1**)^1, 5, 9–11, 22^ revealed dramatic differences in the components of variant interpretations, which were often a mixture of concepts with standardized (e.g. HGNC gene symbols^23^, HGVS variant nomenclature^24^, precise disease terms), externally referenced (e.g. identified elements of an established ontology or database), or knowledgebase-specific (e.g. disease shorthand, internal identifier) representations (**Figure 1**).

**Figure 1.**
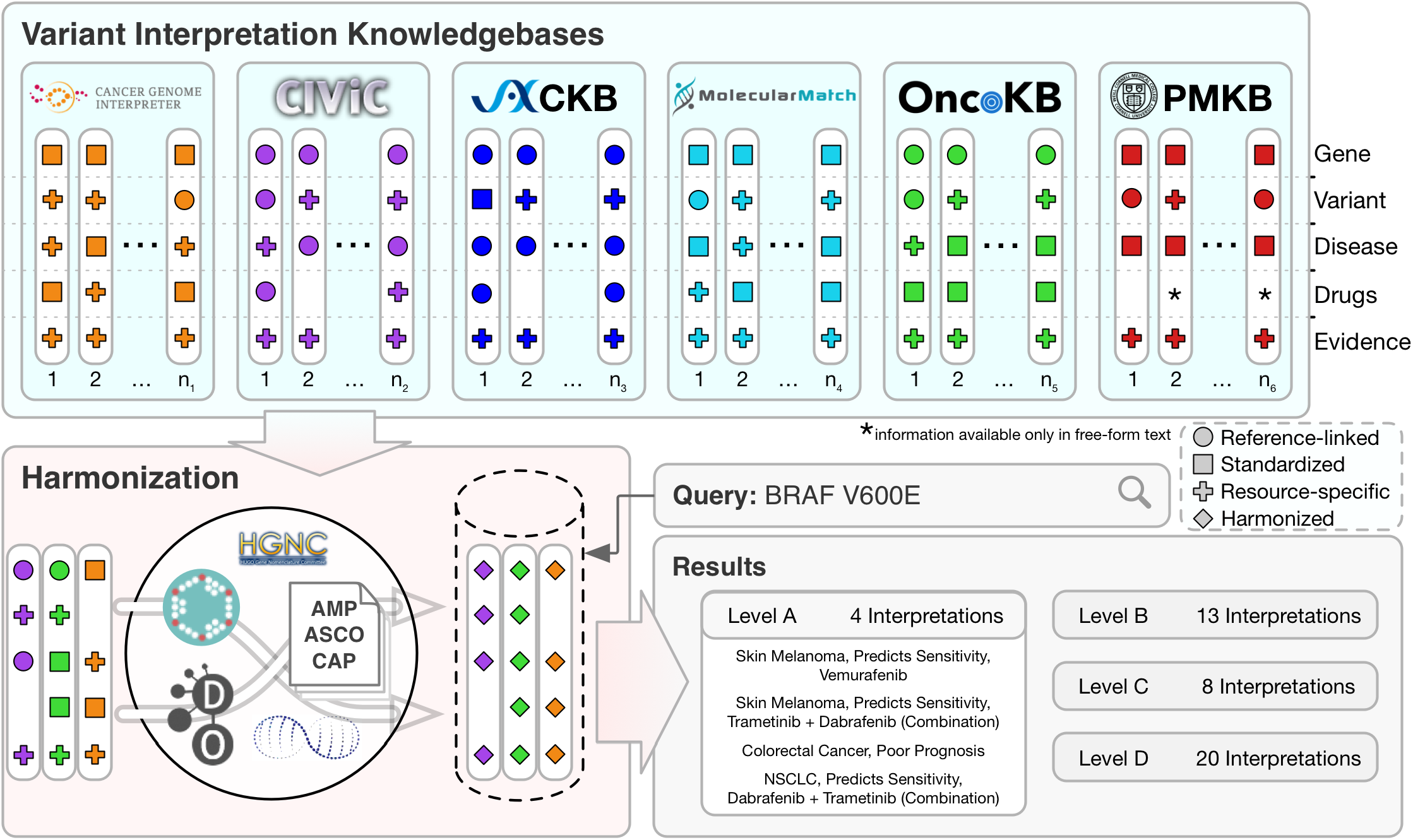
Creation of a harmonized meta-knowledgebase. Six variant interpretation knowledgebases of the VICC (blue panel) and representative symbolic interpretations from each (white columns) are illustrated. Interpretations are split across 5 different elements; **gene, variant, disease, drugs** and **evidence**. Referenced-linked elements correspond to unique identifiers from established authorities for that element (e.g. the use of Entrez or Ensembl gene identifiers). Standardized elements correspond to immediately recognizable formats or descriptions of elements, but are not linked to an authoritative definition. Resource-specific elements are described by terminology unique to the knowledgebase. These elements are each harmonized (red panel) to a common reference standard (shown here is the use of HGNC for genes, ChEMBL for drugs, AMP/ASCO/CAP guidelines for evidence, Disease Ontology for diseases, and ClinGen Allele Registry for variants). This harmonized meta-knowledgebase allows for querying across interpretations from each of the constituent VICC knowledgebases (gray panel, example query *BRAF* V600E), returning aggregated results which are categorized and sorted by evidence level.

To resolve this complexity and provide readily searchable, standardized interpretations across knowledgebases, we evaluated the structure of cancer variant interpretations across the core dataset (**Figure 1**). Our first challenge was to develop a consensus for the minimum required data elements that constitute a cancer variant interpretation. These minimal elements include a gene identifier, variant name, cancer subtype (tumor type and organ), clinical implication (diagnostic, prognostic, therapeutic, or predisposing biomarker), provenance of supporting evidence (e.g., PubMed identifier), and curation source. In addition, we recommended ascribing a tiered level of support for the evidence contributing to the interpretation. Each VICC knowledgebase (**Table S1**) provided cancer variant interpretation knowledge as structured data meeting these requirements.

To adapt disparately-structured interpretations to a common data model, we aggregated cancer variant interpretations from each of these knowledgebases by harvesting their evidence through provider-recommended access methods (e.g. API retrieval, data file downloads). We then harmonized these variant interpretations by mapping all data elements in each knowledgebase to established standards and ontologies describing genes, variants, diseases, and drugs (**Figure 1**). Briefly, genes were harmonized using the Human Gene Nomenclature Committee (HGNC) gene symbol table and include the current HGNC symbol, Ensembl and Entrez gene identifiers. Variants were harmonized through a combination of knowledgebase-specific rules, matching to the Catalog of Somatic Mutations in Cancer (COSMIC)^3^, and use of the ClinGen Allele Registry (reg.clinicalgenome.org). Diseases were harmonized using the European Bioinformatics Institute (EBI) Ontology Lookup Service (OLS; www.ebi.ac.uk/ols/index) to retrieve Disease Ontology terms and identifiers. Drugs were harmonized through queries to the biothings API^25^, PubChem^26^, and ChEMBL^27^, storing the term, description, id and source. Details for each of these harmonization strategies are described in **Online Methods** and **Figure S1**.

Due to the knowledgebase-specific nature of describing an interpretation evidence level (**Figure 1**), harmonization required manual mapping of evidence levels to a common standard. Standards and guidelines for the interpretation and reporting of genomic variants in cancers have been published by the Association for Molecular Pathology (AMP), the American Society of Clinical Oncology (ASCO), and the College of American Pathologists (CAP).^28^ Released after (and partially informed by) the design and curation of the VICC knowledgebases, these guidelines are compatible with (but not identical to) the existing evidence levels of these knowledgebases. We constructed a mapping of evidence levels provided by each knowledgebase to the evidence levels constituting AMP/ASCO/CAP Tier I and II variants (**Table 1**). As a result, variant interpretations can be filtered by Tier I (level A/B) evidence, defined as having strong clinical significance. Interpretations of potential clinical significance (Tier II evidence) comprised of early clinical trials (level C), case studies (level C/D), or preclinical data (level D) are also searchable. Tier III (unknown significance) and Tier IV (benign) variants are not included in this resource.

**Table 1.** Mapping knowledgebase-specific evidence codes to AMP/ASCO/CAP guidelines.

| AMP/ASCO/CAP Variant Evidence Guidelines |  |  |  |  |  |  |  |
| --- | --- | --- | --- | --- | --- | --- | --- |
| Evidence Level | Defining Characteristics | CIViC | OncoKB | JAX-CKB | CGI | MMatch | PMKB |
| Level A<br>(Tier I) | *Evidence from professional guidelines or FDA-approved therapies relating to a biomarker and disease.* |  |  |  |  |  |  |
|  |  | Level A | Level 1 / 2A / R1 | Guideline / FDA Approved | Clinical Practice | Level 1A | Tier 1 |
| Level B<br>(Tier I) | *Evidence from clinical trials or other well-powered studies in clinical populations, with expert consensus.* |  |  |  |  |  |  |
|  |  | Level B | Level 3A | Phase III | Clinical Trials III-IV | Level 1B |  |
| Level C<br>(Tier II) | *Evidence for therapeutic predictive markers from case studies, or other biomarkers from several small studies. Also evidence for biomarker therapeutic predictions for established drugs for different indications.* |  |  |  |  |  |  |
|  |  | Predictive Level C | Level 2B, Level 3B | Clinical Study/ Phase I / Phase II | Clinical Trials I-II, Case Reports | Level 2C | Tier 2 |
| Level D<br>(Tier II) | *Preclinical findings or case studies of prognostic or diagnostic biomarkers. Also includes indirect findings.* |  |  |  |  |  |  |
|  |  | Non-predictive Level C / Level D / Level E | Level 4 | Phase 0, Pre-clinical | Pre-clinical Data | Level 2D | Tier 3 |

Together, these efforts describe a centralized harmonization strategy to structure and unify queries across the knowledgebases describing clinical interpretations of cancer genomic variants (**Figure 1**).

### The landscape of variant interpretation knowledge

The meta-knowledgebase v0.10 release contains 12,856 harmonized interpretations (hereafter referred to as the *core dataset*; **Online Methods**) supported by 4,354 unique publications for an average of 2.95 interpretations / publication. Notably, 87% of all publications were referenced by only one knowledgebase, and only 1 paper^29^ was referenced across all six knowledgebases (**Figure S2a**). Gene symbols were almost universally provided; the few interpretations lacking gene symbols (<0.01%) are structural variants that are not associated with an individual gene. In contrast to publications, the genes curated by the cancer variant interpretation community are much more frequently observed in multiple knowledgebases. We observed that 23% of genes with at least one interpretation were present in at least half of the knowledgebases, compared to only 5% of publications (p < 0.001; Fisher’s exact test; **Figure S2b**).

Variants have little overlap across the core dataset (**Figure 2a**). Of the constituent 3,439 unique variants, 76.6% are described by only one knowledgebase, and <10% are observed in at least three (**Figure 2b**). This lack of overlap is partially due to the complexity of variant representation. For example, the representation of an ERBB2 variant as described in nomenclature defined by the Human Genome Variation Society (HGVS)^24^ is NP_004439.2:p.Y772_A775dup, and yet it is referenced in multiple different forms in the biomedical literature. p.E770delinsEAYVM^30^, p.M774insAYVM^31^, and p.A775_G776insYVMA^32^ all describe an identical protein kinase domain alteration, though they appear to identify different variants (**Figure 2c**). Despite having a standard representation by the HGVS guidelines, these alternative forms continue to appear in the literature, where readers are sometimes explicitly discouraged from the use of the HGVS standard in lieu of historical terms to describe the variant.^32^ Consequently, a researcher looking to identify a specific match to ERBB2 p.E770delinsEAYVM may find no direct matches, though several exist under various alternate representations. This component of variant harmonization is addressed through the use of the ClinGen Allele Registry (**Online Methods**).

**Figure 2.**
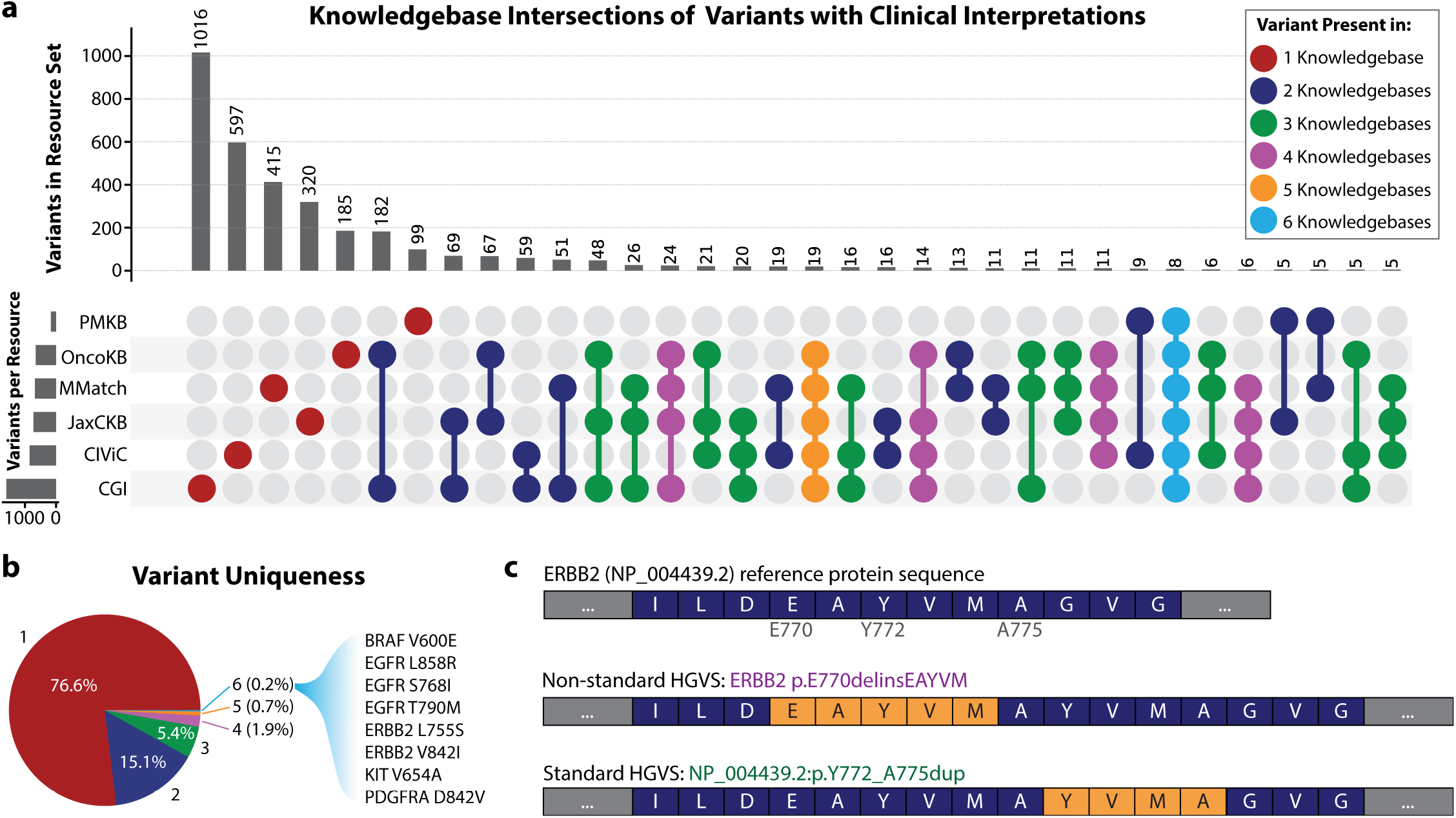
Representation of genomic variants across interpretation knowledgebases. **(a)** UpSet plot^47^ of variants across six cancer variant interpretation knowledgebases. Sets of variant interpretation knowledgebases with shared variants are indicated by colored dots in the lower panel, with color indicating set size (e.g. red dots indicate only the single designated knowledgebase in the set, dark blue dots indicate two knowledgebases in the set, etc.). Objects are attributed to the largest containing set; thus a variant described by all six knowledgebases is attributed to the light blue set with 8 variants. **(b)** Pie chart visualizing overall uniqueness of variants, with categories indicating the number of knowledgebases describing each variant. Nearly 77% of variants are unique across the knowledgebases, with only 0.2% ubiquitously represented. The 8 variants present in all 6 knowledgebases are listed at right. **(c)** Multiple syntactically-valid representations of an identical protein product can lead to confusion in describing the change in the literature and in variant databases. At top, the wild-type protein sequence is represented for ERBB2. Below, two (of many) possible representations of an in-frame insertion are shown. In the middle a non-standard HGVS expression describes a 5 amino acid insertion. At the bottom, the HGVS standard representation shows an identical protein product from a 4 amino acid duplication. A search for one representation against a database with another (non-overlapping) representation may lead to omission of a clinically relevant finding. PMKB=Precision Medicine Knowledgebase, CIViC=Clinical Interpretations of Variants in Cancer, CGI=Cancer Genome Interpreter, JAX-CKB=The Jackson Laboratory Clinical Knowledgebase, MMatch=MolecularMatch interpretation knowledgebase.

To illustrate this, we performed a survey of all interpretations describing the previously discussed ERBB2 variant (NP_004439.2:p.Y772_A775dup) using the public web search interface provided by each knowledgebase (**Tables 2, S2**). Each knowledgebase that had an entry for this variant represented it differently. Two did not have specific interpretations for this variant, though they did have relevant container mutations (e.g., *exon 20 insertions*; **Table 2**). Most of the knowledgebases had a single internal representation of the variant, although the majority of these terms did not match across knowledgebases. The evidence describing these interpretations varied considerably in form, as each used knowledgebase-specific nomenclature (e.g. evidence described as “Level 3A” in OncoKB is equivalent to “Level 1B” from MolecularMatch, or “Level B” from CIViC; **Tables 1, 2**). Of the 19 unique publications describing the collected evidence, only 3 (two American Association for Cancer Research [AACR] abstracts and one journal article) were observed in more than one knowledgebase, and none were observed in more than two. Interestingly, the curated interpretations from these shared publications varied by knowledgebase in disease scope (*advanced solid tumor* compared to *non-small cell lung cancer^33^*; *breast cancer and non-small cell lung cancer* compared to *cancer^34^*). A review of the interpretations revealed some that are present in most of the knowledgebases (e.g. *use of afatinib, trastuzumab, or neratinib in non-small cell lung carcinomas*; **Table 2**), and others that are present in only one or two (e.g. *use of lapatinib in lung adenocarcinoma* and *use of afatinib and rapamycin in combination* are observed in only one knowledgebase each; **Table 2**). Importantly, this includes sparse interpretations that describe conflicting evidence (e.g. *no benefit from neratinib in non-small cell lung carcinoma*; **Table 2**) or negative evidence (e.g. *does not support sensitivity/response to dacomitinib in NSCLC*; **Table 2**). Collectively, these data illustrate the diversity in knowledgebase structure, content, terminology, and curation methodology. Consequently, utilizing subsets (or alternate sets) of knowledgebases will very likely result in differing sets of interpretations.

**Table 2.** Comprehensive assessment of NP_004439.2:p.Y772_A775dup variant across clinical interpretation knowledgebases.

| Resource | ERBB2 Variant | Evidence | Document ID | Interpretation |
| --- | --- | --- | --- | --- |
| CIViC | M774INSAYVM | Level B, 2-star | PMID:25899785 | Does not support sensitivity/response to Dacomitinib in NSCLC |
|  | M774INSAYVM | Level C, 4-star | PMID:26559459 | Supports sensitivity/response to Afatinib in Lung Adenocarcinoma |
|  | M774INSAYVM | Level C, 3-star | PMID:22325357 | Supports sensitivity/response to Afatinib in Lung Adenocarcinoma |
|  | M774INSAYVM | Level C, 3-star | PMID:25789838 | Supports sensitivity/response to Trastuzumab Emtansine in Lung Adenocarcinoma |
|  | M774INSAYVM | Level D, 3-star | PMID:19122144 | Supports sensitivity/response to Afatinib and Rapamycin (combination) in NSCLC |
|  | Kinase Domain Mutation | Level C, 4-star | PMID:26598547 | Supports sensitivity/response to Trastuzumab in Lung Adenocarcinoma |
|  | Kinase Domain Mutation | Level C, 3-star | PMID:22325357 | Supports sensitivity/response to Afatinib in Lung Adenocarcinoma |
| OncoKB | Exon 20 Insertions | Level 4 | 10.1158/1538-7445.AM2016-2644 | Supports response to AP32788 in Non-Small Cell Lung Cancer |
|  | Oncogenic Mutations | Level 3A | PMID:23220880<br>10.1158/1538-7445.AM2017-CT001 | Supports response to Neratinib in Breast Cancer and Non-Small Cell Lung Cancer |
| CGI | inframe ins. A775YVMA | Early trials | 10.1200/JCO.2017.35.15_suppl.8510 | Responsive to Ado-Trastuzumab Emtansine in Lung Cancer |
|  | inframe ins. A775YVMA | Early trials | 10.1158/1538-7445.AM2017-CT001 | Responsive to Neratinib in Cancer |
|  | proximal exon 20 | Early trials | PMID:26598547 | Responsive to Afatinib, Neratinib, Lapatinib, or Trastuzumab in Lung Adenocarcinoma |
|  |  |  | 10.1200/JCO.2017.35.15_suppl.9071 |  |
| PMKB | exon(s) 20 insertion | Tier 2 | PMID:22761469 | Associated with sensitivity to some ERBB2 inhibitors in Lung Adenocarcinoma |
|  |  |  | PMID:16818618 |  |
|  |  |  | PMID:25152623 |  |
| JAX-CKB | Y772_A775dup | Clinical Study | PMID:26964772 | Conflicting response to afatinib in Lung Adenocarcinoma |
|  | Y772_A775dup | Phase II | PMID:29420467 | Predicted sensitive to neratinib in Her2-receptor negative breast cancer |
|  | Y772_A775dup | Phase II | PMID:29420467 | Predicted resistant to neratinib in urinary bladder cancer and non-small cell lung carcinoma |
|  | Y772_A775dup | Preclinical | PMID:26545934 | Sensitive to afatinib in lung cancer |
|  | Y772_A775dup | Preclinical | PMID:26545934 | No benefit to gefitinib in lung cancer |
|  | Y772_A775dup | Preclinical | PMID:28363995 | Sensitive to neratinib in advanced solid tumor |
|  | exon 20 insertion | Clinical Study | PMID:28167203 | Predicted sensitive to afatinib or trastuzumab in non-small cell lung carcinoma |
|  | exon 20 insertion | Clinical Study | PMID:26964772 | Predicted sensitive to afatinib in lung adenocarcinoma |
|  | exon 20 insertion | Phase II | PMID:29420467 | Predicted sensitive to neratinib in Her2-receptor negative breast cancer |
|  | exon 20 insertion | Phase II | PMID:29420467 | No benefit to neratinib in non-small cell lung carcinoma |
|  | exon 20 insertion | Preclinical | 10.1158/1538-7445.AM2016-2644 | Sensitive to AP32788 in advanced solid tumor |
| Molecular Match | Y772_A775dup | Level 1B | PMID:22325357, 26964772 | Confers sensitivity to Afatinib in patients with Neoplasm of lung |
|  | Y772_A775dup | Level 2C | PMID:26598547 | Confers sensitivity to Trastuzumab in patients with Neoplasm of lung |
|  | Y772_A775dup | Level 2D | PMID:22325357 | Confers sensitivity to Afatinib in patients with Neoplasm of breast |
|  | A775_G776insYVMA | Level 1A | PMID:26559459, 22325357, 26545934 | Confers sensitivity to Afatinib in patients with Neoplasm of lung |
|  | A775_G776insYVMA | Level 2C | PMID:23610105, 26964772, 22908275 | Confers sensitivity to Afatinib in patients with Neoplasm of breast |
|  | A775_G776insYVMA | Level 2D | PMID:17311002, 22908275 | Confers sensitivity to Neratinib in patients with Neoplasm of breast |

### Harmonization improves consensus across interpretations

To test the effect of our harmonization methods on generating consensus, we evaluated the overlap of unique interpretation elements from each knowledgebase of the core dataset in comparison to evaluated unharmonized (but aggregated) data (**Online Methods**). As noted above, genes from each resource used HGNC gene symbols, resulting in very little gain from harmonization; 45% of genes across knowledgebases overlapped without harmonization, compared to 46% with harmonization. This is in contrast to variants (8% overlapping unharmonized, 26% overlapping harmonized), diseases (27% unharmonized, 34% harmonized), and drugs (20% unharmonized, 36% harmonized) (**Table S3)**. None of the evidence levels were consistent across resources when unharmonized, and all are consistent with a common standard (**Table 1**) after harmonization, a primary contribution of this work.

### Harmonization of variant interpretations increases findings of strong clinical significance

Evaluation of patient variants for strong clinical significance requires an assessment of these variants in the appropriate disease context. The aggregated knowledge across the core dataset describes 357 distinct disease concepts from the Disease Ontology (DO)^35^ across 12,497 interpretations (**Table S4**). These diseases range from highly specific (e.g. *DOID:0080164 - myeloid and lymphoid neoplasms with eosinophilia and abnormalities of PDGFRA, PDGFRB, and FGFR1*) to generalized (e.g. *DOID:162 - cancer*). To compare the variant interpretations to disease type, we used the expert-curated “TopNodeCancerSlim” DO mapping^36^ that describes 58 common, top-level disease terms (TopNode terms) across several major datasets, including The Cancer Genome Atlas (TCGA), International Cancer Genome Consortium (ICGC), and COSMIC.^3, 37, 38^ When linked to the nearest TopNode term, 5 major cancer group terms each accounted for over 5% of all interpretations in the core dataset: *lung cancer* (24%), *breast cancer* (13%), *hematologic cancer* (11%), *large intestine cancer* (9%), and *melanoma* (6%) (**Figure 3a** and **Table S5**). Notably, the most common interpretations mirror TopNode terms that have both high incidence (**Figure 3b**) and high mortality (**Figure 3c**) as reported by the National Cancer Institute (**Table S6**)^39^: *lung cancer*, *breast cancer*, and *hematologic cancer*. The *large intestine cancer* TopNode term contains numerous interpretations describing *colorectal cancers*, which are highly applicable to the related TopNode *colon cancer* (a top-five cancer in both incidence and mortality; **Table S7**). Evaluation of these terms across the core dataset revealed significant differences in the distribution of common cancer types constituting each knowledgebase, illustrating the value of aggregating knowledgebases for a more comprehensive landscape of interpretations (**Figure S3**, **Table S8**).

**Figure 3.**
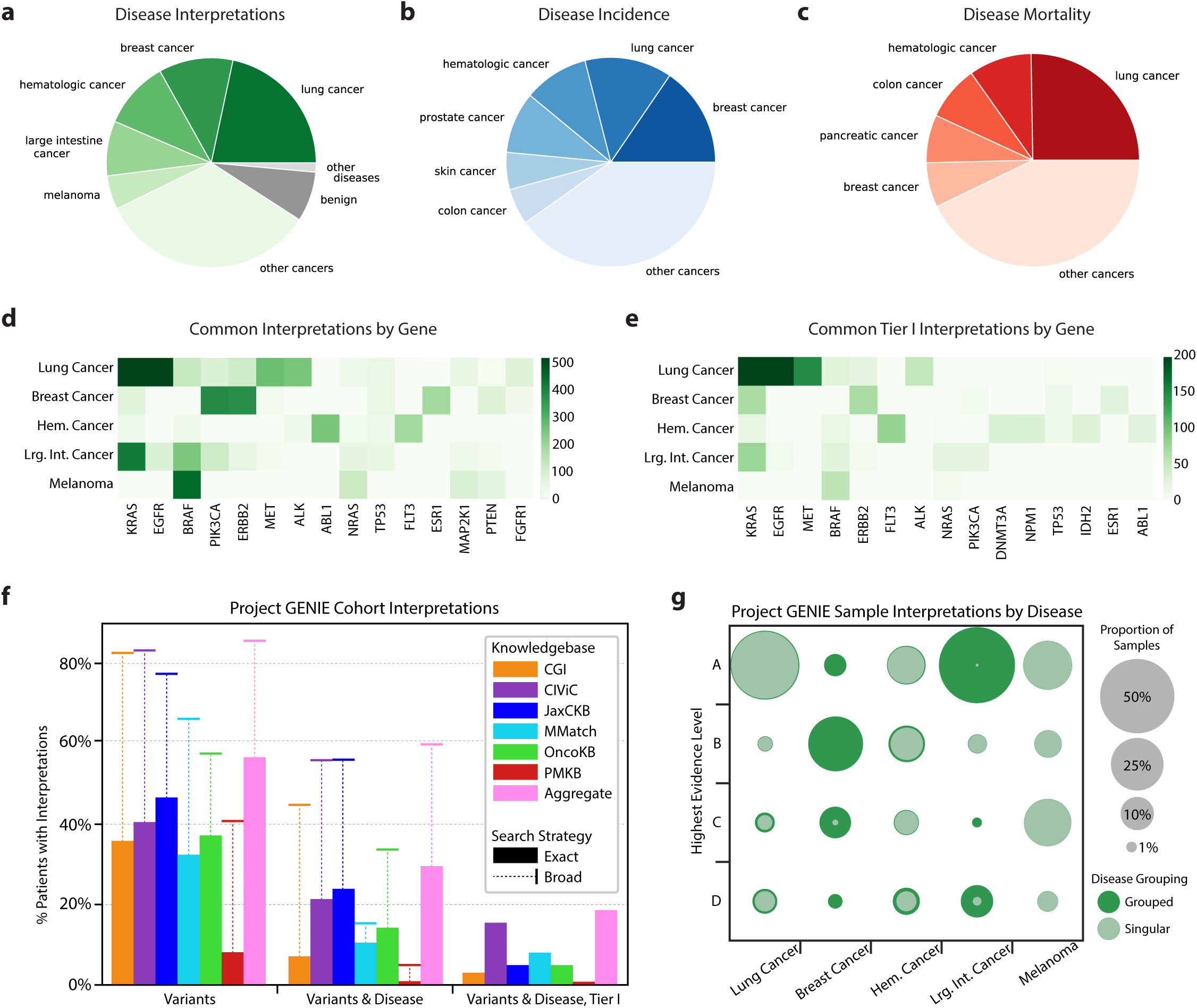
Clinical interpretations of variants are defined by disease. **(a)** Core dataset interpretations for top-level disease groups. Distinct diseases are shown if the constituent interpretations for that disease account for at least 5% of the total dataset. Diseases accounting for at least 5% of cancer incidence **(b)** and mortality **(c)** are also displayed. Approximately 8% of interpretations are categorized as benign neoplasms (dark gray; e.g. von Hippel-Lindau disease). An additional 1% are categorized under high-level terms other than *DOID:14566 - Disease of Cellular Proliferation*. **(d)** A heatmap of frequent gene-disease interpretations, and **(e)** the related heatmap limited to tier 1 interpretations. **(f)** Percentage of Project GENIE cohort with at least one interpretation from the indicated knowledgebase that matches patient variants (left group), patient variants and disease (center group), or patient variants, disease, and a Tier I evidence level (right group). A broader search strategy (indicated by whisker bars; **Figure S4**) that allows for regional variant matches (e.g. gene-level) and broader interpretation disease terms (e.g. *DOID:162 - cancer*) nearly doubles the number of patients with matching interpretations. These broader match strategies are incompatible with the ASCO/AMP/CAP evidence guidelines. **(g)** Most significant finding (by evidence level) across patient samples, by disease. Each column represents one of the common diseases indicated in **(a)**, and the rows represent the evidence levels described in **Table 1**. Inner, light green circles (labeled *Singular*) indicate the proportion observed when matching patient diseases to interpretations with the same disease ontology term. Outer, dark green circles (labeled *Grouped*) indicate the proportion observed when matching patients to interpretations with ancestor or descendant terms that group to the same class of disease (**Online Methods**). Hem. Cancer=Hematological Cancer, Lrg. Int. Cancer=Large Intestine Cancer.

To further test the value of harmonized interpretation knowledge, we evaluated the 38,207 patients of the AACR Project Genomics Evidence Neoplasia Information Exchange (GENIE)^40^. We first queried the 237,175 moderate or high impact variants from GENIE using a broad search strategy (**Online Methods** and **Figure S4a**). Notably, 11% (4,305) of patients lacked any variants to search before filtering on predicted impact, and 12% (4,355) after. This search yielded 2,316,305 interpretation search results for an average of 9.8 interpretations / variant query. For a point mutation, these interpretations included matches to alternate alleles at the same position, associated amino acid changes, the exon or functional domain, or gene-level interpretations such as *overexpression*, *gain/loss-of-function*, or simply *mutations*. Restricting to an exact coordinate match (and thus excluding gene-level interpretations; **Figure S4a** *positional match*) revealed an interpretation result set dominated by a few common GENIE point mutations in variants each with a large number of interpretations, including BRAF NP_004324.2:p.V600E, KRAS NP_004976.2:p.G12 mutations, and both NP_006209.2:p.E545K and NP_006209.2:p.H1047R mutations in PIK3CA (**Figure S5**). This is congruent with our observation that the interpretations of the core dataset for the most common diseases are highly focused on these and other specific genes (**Figure 3d**), including Tier I interpretations (**Figure 3e**). Examining our results at the patient level revealed that a focused, variant-level search resulted in at least one interpretation (in any cancer type with any level of evidence) for 57% of patients in the GENIE cohort, compared to the average 33% obtained when using each constituent knowledgebase individually (**Figure 3f**). We observed that broadening the search scope to include any overlapping variants (see **Figure S4a** *regional match*) increased the cohort coverage to 86% of patients (compared to an average of 68% per individual knowledgebase). However, it is prudent to keep in mind that the increase in matching percentage using regional match instead of exact match would be partly due to non-oncogenic passenger variants.

A key component in determining the clinical relevance of an interpretation is whether the tumor type reported in the interpretation matches the patient’s tumor type (See *Defining Characteristics*, **Table 1**). To evaluate this question, we first mapped the tumor types of the GENIE cohort to DO terms. GENIE samples are annotated with a diverse array of Oncotree ontology (oncotree.mskcc.org) disease codes, with 81% (539 / 667) of Oncotree diseases represented in the dataset. Over 55% (299 / 539) of the Oncotree diseases from GENIE do not link to DO through cross-references, of which 41% (123 / 299) do not have any cross-references (**Table S9**). This lack of cross-references among GENIE diseases is significantly higher than the 25% (166 / 667) of all Oncotree terms lacking cross-references (p < 0.001; Fisher’s exact test), suggesting that terms used to describe individual patient cancers (e.g. *Well-Differentiated Neuroendocrine Tumor of the Rectum*) are less likely to map to other knowledgebases than high-level parent terms (e.g. *colorectal cancer*). Despite this, 80% of GENIE patients had a disease term map to DO, indicating that the common cancers among this cohort are more likely to be cross-referenced adequately for mapping. Further evaluation confirmed a significant enrichment of more frequently observed disease terms among the terms that mapped to DO, compared to those that did not (p = 0.002; Mann-Whitney U test). Restricting patient search results to those interpretations that are of matched grouped (TopNode) disease terms (**Figure S4b**; **Online Methods**) resulted in 29% of patients with at least one clinical interpretation (compared to an average individual knowledgebase match rate of 13%), and 18% of patients with at least one Tier I clinical interpretation (compared to an average 6% per individual knowledgebase) (**Figure 3f**). Allowing matching to any ancestor or descendant term and allowing partial variant overlaps improves matches to 60% (compared to an average of 35% per individual knowledgebase). This broader strategy, however, requires contextual re-evaluation of assigned AMP/ASCO/CAP evidence levels, which are designated for a precise match to variant and disease context. Consequently, evidence level or tier filtering can only be used with an exact search strategy. An even broader search strategy (**Figure S6**) that allows variant matching to interpretation genes has comparable findings to the overlapping variant strategy, indicating that many of the commonly mutated genes have gene-level interpretations.

A comparison of interpretations across the identified common cancers revealed that the use of grouped terms instead of exact terms for matching interpretations to patients’ cancers varies dramatically by cancer type, with some cancers (e.g. *lung cancer, melanoma*) showing little increased interpretation breadth, while others have enormous effect (e.g. *breast cancer*, *large intestine cancer*; **Figure 3g**). This is primarily due to the specific nature by which patients are classified with certain diseases, versus the aggregate nature by which interpretations are ascribed to diseases. Interestingly, 50% of GENIE patient samples have disease-matched interpretations across the frequently observed cancers, compared to only 33% of patient samples across all other cancers (p < 0.001; Fisher’s exact test). These numbers are reduced to 39% and 15%, respectively, when considering only Tier I interpretations (p < 0.001; Fisher’s exact test).

### A resource for searching aggregated and harmonized variant interpretation knowledge

We have developed and hosted a public web interface for exploring the VICC meta-knowledgebase, freely available online at search.cancervariants.org. This interface accesses an ElasticSearch index for the most recent data release of the VICC harmonized knowledgebase. Searching the knowledgebase is performed through specifying filters for any term or entering free text or compound (e.g. and/or logic) queries in the search box at the top of the page (**Figure 4a**). Panels with data distribution visualizations describe the current result set (**Figure 4b**). These interactive panels provide additional information about specific subsets, and may be used to create additional filters (e.g. clicking on a level in the *evidence_level* panel filters results throughout the page to display only those interpretations with the designated evidence level). This allows investigators to see the distribution of interpretations by evidence level, disease, gene, and drug, and filter according to their interests. Tabulated results are provided at the bottom of the page (**Figure 4c**), and are expandable with all details, including the (unharmonized) record provided by the original knowledgebase for each interpretation. These search tools are available via both the web interface and an API search endpoint (**Online Methods**), in addition to a GA4GH beacon on beacon-network.org. Additionally, a Python interface and analysis workbook have been developed to enable reproduction (and additional exploration) of the data presented in this paper, as well as full downloads of the underlying data (**Online Methods**).

**Figure 4.**
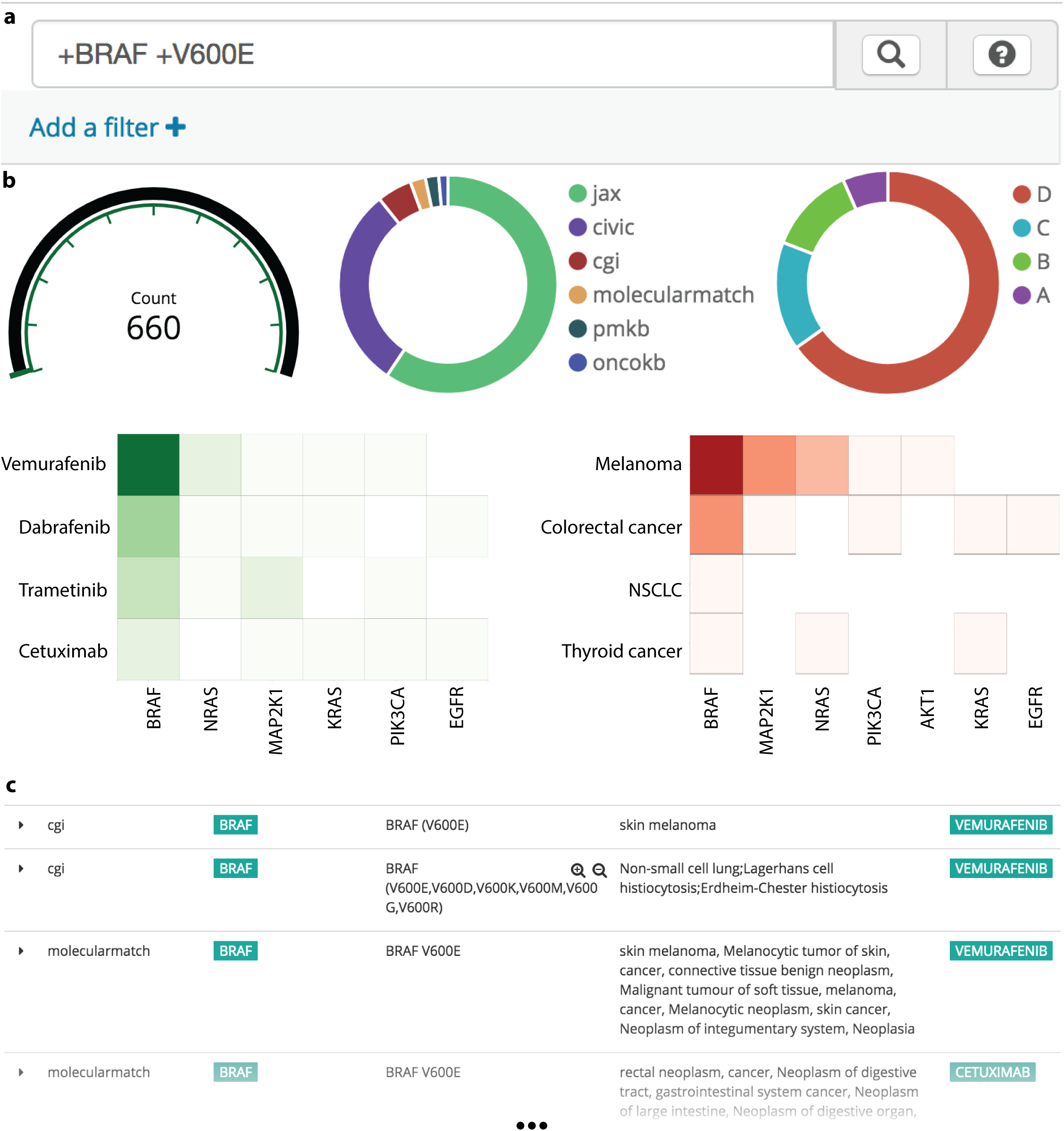
A web client for exploring the VICC meta-knowledgebase. **(a)** Queries are entered as individual terms, with compound queries (e.g. *BRAF* and *V600E*) denoted by preceding ‘+’ characters. Usage help and example documentation can be found by clicking the “?” icon. **(b)** Result visualization panels are interactive, allowing users to quickly filter results by evidence level, source, disease, drug, and gene. **(c)** Scrollable results table has sortable columns detailing each resource (e.g., *molecularmatch*), gene (*BRAF*), variant (*V600E*), disease (*skin melanoma*), drug (*vemurafenib*), evidence level, evidence direction, original URL, and primary literature. Rows are expandable and include additional detail structured as both JSON and a table.

## DISCUSSION

In this study, we aggregated and analyzed clinical interpretations of cancer variants from six major knowledgebases^1, 5, 9–11, 22^. Our analysis uncovered highly disparate content in curated knowledge, structure, and primary literature across these knowledgebases. Specifically, we evaluated the unique nature of the vast majority of genomic variants reported across these knowledgebases, and demonstrated the challenge of developing a consensus interpretation given these disparities. These challenges are exacerbated by non-standard representations of clinical interpretations, in both the primary literature and curated knowledge of these resources. It is encouraging that the curators of these knowledgebases have, without coordination, independently curated diverse literature and knowledge sources. However, this reflects an enormous curation burden generated from the increasingly common molecular characterisations of patient tumors and the related expansion of primary literature describing them. Our findings thus highlight the need for a cooperative, global effort to curate comprehensive and thorough clinical interpretations of molecular variants for robust practice of precision medicine.

We formed the Variant Interpretation for Cancer Consortium (VICC) of leaders of prominent cancer variant knowledgebases and experts in variant interpretation, software development, curation, ontologies, and clinical translation to implement a framework to address the challenge of aggregating and cohesively representing variant interpretation knowledge. In doing so, we first defined the distinct elements of a clinical interpretation (gene, variant, disease, drug, evidence), and then extracted these data from each of the constituent knowledgebases hosted by institutions of the VICC. Extracted data were harmonized to established reference standards for each element and stored in a centralized database. We developed a prototype web interface, API, and python package for accessing and querying these data. Together, these tools provide the foundation to analyze the aggregate interpretation knowledge across the constituent knowledgebases, and are freely available and open source (MIT-licensed; see **Online Methods**) for public use. The content of the meta-knowledgebase is dynamic, as we routinely poll the constituent knowledgebases for the current clinical interpretations of cancer variants.

Harmonization improved concordance between interpretation elements across resources, with large gains in overlapping terms between resources across variants, diseases, and drugs. Importantly, this harmonization allowed us to search relationships between patient and interpretation disease terms, improving precise matching between patient and interpretations of clinical significance. We noted that there was little need to harmonize gene identifiers, as each knowledgebase had independently selected HGNC gene symbols as a reference, enabling easy and direct comparison of genes. This underscores both the utility of standardized data, as well as the need for adoption of similar standards (such as those described in this work) to drive direct comparison of variant interpretations. In our analysis of the variants and diseases of the harmonized interpretations, we observed that frequent top-level cancer terms mirror cancers with high incidence and mortality. We also noted that a large percentage of these interpretations described a relatively small number of gene-disease relationships. We subsequently searched the meta-knowledgebase for interpretations describing the patients of the GENIE cohort. As a result of our harmonization of interpretations across knowledgebases, we were able to achieve at least one specific (position-matched) variant interpretation for 57% of the patients in the cohort. In the most stringent searches we required a precise variant match to a Tier I interpretation also matching the patient’s cancer; in these cases, 18% of the cohort had a finding of strong clinical significance. Notably, these findings were substantially higher in patients with more common cancers, with 39% of the common cancer samples variant-matching at least one Tier I interpretation, compared to 15% of other cancer samples. These findings are concordant with observations of matched therapy rates in precision oncology trials, including 15% from IMPACT/COMPACT^15^, 11% from MSK-IMPACT^14^, 5% from the MD Anderson Precision Medicine Study^16^ and 23% from the NCI-MATCH trials^17^.

Collectively, our results portray a confluence of knowledge describing the most common genomic events relevant to the most frequent cancers, with highly disparate knowledge describing less frequent events in rare cancer types. The differing content of these knowledgebases may be a result of research programs targeted at frequent cancers, highlighting a need for a broader focus on less common cancers. This sparse landscape of curated interpretation knowledge is exacerbated by paucity in cross-references between ontologies describing disease, highlighting the importance of bridging this gap^41^. Similarly, complexities in variant representation have elucidated a need for sophisticated methods to harmonize genomic variants; harmonization with the ClinGen Allele Registry (reg.clinicalgenome.org) is suited to point mutations and indels, but the representation and harmonization of complex and non-genomic (e.g. expression, epigenetic) variants remains a challenge.

Our harmonized clinical interpretation meta-knowledgebase represents a significant step forward in building consensus that was previously unattainable due to a lack of harmonization services such as the Allele Registry and expert standards and guidelines such as those recommended by AMP/ASCO/CAP. This meta-knowledgebase serves as an open resource for evaluating interpretations from institutions with distinct curation structure, procedures, and objectives. Potential uses include expert-guided therapy matching, supporting FDA regulatory processes associated with lab-developed genomic tests for guiding therapy, and identification of diseases and biomarkers that warrant future study.

While our initial efforts provide a structure by which variant interpretation knowledgebases can contribute to a broader and more consistent set of interpretations, much work remains to be done. In particular, VICC members contribute to GA4GH Work Streams to develop and integrate new and existing^42–45^ standards for the representation of variant interpretations and the evidence that describe them. Our web interface is being redesigned to a full-scale web service and user interface to concisely represent the most relevant interpretations for one or more variants. Additionally, we are building inference tools to automatically identify the concepts users are querying in real time. We also will be expanding our effort to harmonize and present interpretations of various non-coding variants, structural variants beyond gene pairs, and aggregate markers like microsatellite instability status. A prioritized long-term goal is the development of standards and techniques for interpretations of combined germline and somatic variations. Similarly, we are building guidelines and methods to enable automated consensus recommendations. Finally, we are seeking out additional knowledgebases of clinical interpretations of variants to harmonize and share with the broader cancer genomics community, and building an API specification which they may use to incorporate their own interpretations.

In conclusion, there is a great need for a collaborative effort across institutions to build structured, harmonized representations of clinical interpretations of cancer genomic variants to advance precision medicine implementation. Our work has illustrated the diversity of variant interpretations available across resources, leading to inconsistency in interpretation of cancer variants. We have assembled a framework and recommendations for structuring and harmonizing such interpretations, from which the cancer genomics community can improve consensus interpretation for cancer patients. Anyone can leverage the open and freely available aggregated knowledge resources, and associated software tools, described in this work at search.cancervariants.org. Our working group and open source software development environment are open to all and we welcome participation from anyone with interest in learning about, utilizing, augmenting, improving, or proposing new directions for this community-based project, for the benefit of cancer patients.

## ACKNOWLEDGEMENTS

We would like to acknowledge the contributions from members of GA4GH and specifically the Genotype to Phenotype Task Team for their numerous contributions leading to this study. We would like to acknowledge the VICC for their input in construction of the meta-knowledgebase and drafting of the paper. We would like to acknowledge Matthew McCoy for his assistance in proofreading the manuscript. AHW was supported by a fellowship from the NCI (NIH Grant F32CA206247). MH was supported by the Monarch Initiative NIH Office of Director 5R24OD011883. JiG, DC, and NS were supported by a National Cancer Institute Cancer Center Core Grant (P30-CA008748). ML was supported through the Medical Research Council - Cancer Research UK Stratification in Colorectal Cancer Program grant, and Health Data Research UK Substantive Site grant. MG was supported by a career development award from the NHGRI (NIH Grant R00HG007940). OLG, MG and the CIViC knowledgebase were supported by the NCI (NIH Grant U01CA209936).

## AUTHOR CONTRIBUTIONS

BW, GM, JeG, and AHW developed the harvester and normalization routines. DT, JDP, and NL guided harvesting content from CGI. AHW, KK, OLG, and MG guided harvesting content from CIViC. SP and SM guided harvesting from JAX-CKB. RPD and XSL guided harvesting content from MolecularMatch. JiG and DC guided harvesting content from OncoKB. OE guided harvesting content from PMKB. RD provided case studies illustrating need for harmonization. DS introduced strategy for harmonizing evidence level, and AHW mapped the evidence levels with feedback from DT, NL, KK, OLG, MG, SM, XSL, DC, and OE. LMS, JM and MH contributed discussion of harmonizing ontologies. AHW developed the python interface to the dataset. AHW and KK created Table S1. KK created Table S8 and Figure S3. AHW created all other tables and figures. BW developed the prototype web client and API. AAM led the genotype to phenotype working group and informed the harmonization strategy. OLG, MG, NL and DT founded and led the VICC. OLG, MG, AAM, and JeG supervised the project. AHW wrote the manuscript, with regular feedback from all authors. All authors contributed in weekly discussions and in revising and approving the manuscript.

## ONLINE METHODS

### Harvesting cancer variant interpretation knowledge

OncoKB, the Cancer Genome Interpreter (CGI) and JAX-Clinical Knowledgebase (JAX-CKB) all contain complementary knowledge of variant oncogenicity. While valuable, knowledge of a variant’s potential role in driving tumorigenesis is structured differently than clinical interpretations of genomic variants, and is therefore outside of the scope of this manuscript. While omitted from the analyses presented in this paper, we do aggregate these annotations due to their potential utility in clinical research. ClinGen, ACMG, AMP, ASCO, and CAP are working on developing guidelines in order to enable consistent and comprehensive assessment of oncogenicity of somatic variants. In the future, variant oncogenicity interpretations based on such guidelines can be incorporated into meta-knowledgebase and should help to improve the harmonization of related interpretations.

Exact code for harvesting and harmonizing each of the VICC knowledgebases may be found online at https://github.com/ohsu-comp-bio/g2p-aggregator. The cancer biomarker database from CGI was harvested from the cgi_biomarkers_per_variant.tsv file from the biomarkers download at https://www.cancergenomeinterpreter.org/data/cgi_biomarkers_latest.zip. CIViC content was harvested via the gene and variant API endpoints documented online at http://griffithlab.org/civic-api-docs/. JAX-CKB content of the publically available 86 genes were harvested from an unpublished API endpoint (harvester code online at https://github.com/ohsu-comp-bio/g2p-aggregator/blob/v0.10/harvester/jax.py#L145-L147). MolecularMatch content was harvested via an authorized API key for use in the aggregated knowledgebase (harvester code online at https://github.com/ohsu-comp-bio/g2p-aggregator/blob/v0.10/harvester/molecularmatch.py). OncoKB content was harvested via a combination of the levels, genes, variants, and variants/lookup API endpoints documented online at: http://oncokb.org/#/dataAccess. PMKB content was provided as a JSON file by the knowledgebase, which we are hosting online at: https://s3-us-west-2.amazonaws.com/g2p-0.7/unprocessed-files/pmkb_interpretations.json

### Harmonizing genes

Gene symbols were matched to the table of gene symbols from HGNC, hosted at the European Bioinformatics Institute (EBI)^46^: ftp://ftp.ebi.ac.uk/pub/databases/genenames/new/json/non_alt_loci_set.json. This table was used to construct an “Aliases” table comprised of retired and alternate symbols for secondary lookup if the interpretation gene symbol was not found among the primary gene symbols from HGNC. If an alias used by a knowledgebase was shared between two genes, omitted by the knowledgebase, or failed to match either the primary or alias table, the gene was omitted from the normalized gene field.

### Harmonizing variants

Variants harvested from each knowledgebase were first evaluated for attributes specifying a precise genomic location, such as chromosome, start and end coordinates, variant allele, and an identifiable reference sequence. Variant names were queried against the Catalog of Somatic Mutations in Cancer (COSMIC)^3^ v81 to infer these attributes in knowledgebases that did not provide them. Custom rules were written to transform some types of variants without clear coordinates (e.g. amplifications) into gene coordinates. All variants were then assembled into HGVS strings and submitted to the ClinGen Allele Registry (http://reg.clinicalgenome.org) to obtain distinct, cross-assembly allele identifiers, if available.

### Harmonizing diseases

Diseases were matched to the Disease Ontology (DO),^35^ through lookup with the European Bioinformatics Institute (EBI) Ontology Lookup Service (OLS)^46^, unless a pre-existing ontology term for a different ontology existed (98.7% of interpretations map to DO). We downloaded the March 2018 release of the TopNode terms from https://github.com/DiseaseOntology/HumanDiseaseOntology/blob/master/src/ontology/subsets/TopNodes_DOcancerslim.json and mapped our interpretation diseases to this list, assigning each disease to its nearest TopNode ancestor (**Table S4**). We assigned remaining interpretation diseases to the non-specific term of *DOID:162* - *Cancer* if the disease was a descendent of this term, but not a descendant of one of the TopNode terms.

### Harmonizing drugs

Drug names were first queried against the biothings API^25^ for harmonization (http://c.biothings.io/v1/query) and if not found were subsequently queried against the PubChem Compounds (https://pubchem.ncbi.nlm.nih.gov/rest/pug/compound/)^26^, PubChem Substances (https://pubchem.ncbi.nlm.nih.gov/rest/pug/substance/), and ChEMBL (https://www.ebi.ac.uk/chembl/api/data/chembl_id_lookup/search)^27^ web services.

### Harmonizing evidence level

Evidence levels were standardized to the AMP/ASCO/CAP guidelines as outlined in **Table 1**.

### Comprehensive evaluation of ERBB2 duplication

Public web portals for the six VICC knowledgebases were manually searched for interpretations for variants describing the alteration detailed in **Figure 2c**. The web portals are freely available online without registration at the following URLs:

- CGI: https://www.cancergenomeinterpreter.org/biomarkers
- CIViC: https://civicdb.org/search/variants/
- JAX-CKB: https://ckb.jax.org/geneVariant/find
- MolecularMatch: https://app.molecularmatch.com/
- OncoKB: http://oncokb.org
- PMKB: https://pmkb.weill.cornell.edu

### Evaluating non-harmonized aggregate content

To evaluate the gains provided by our harmonization methods, we collected and minimally formatted interpretation elements from each knowledgebase without using any harmonization routines. We selected the set of unique elements for each resource and calculated the overlap across the union of those sets (**Table S3**). We then repeated this procedure for harmonized elements and compared total element count and percent overlap between harmonized and non-harmonized elements.

For **genes**, we used HGNC gene symbols, which were provided by each knowledgebase. Gene symbols were almost universally provided across interpretations, although some interpretations do not have associated genes.

For **variants**, we extracted the genomic coordinates (chromosome, start, stop) from each resource and created a unique set of those variants. JAX-CKB and OncoKB do not provide genomic coordinates for variants. When applicable, we split records by the appropriate delimiter to separate out multiple variants. For CGI, we also did minimal HGVS parsing for chr/start/stop when gDNA HGVS strings were provided.

For **diseases**, we extracted the disease term from each knowledgebase and transformed it to lowercase text. PMKB represents diseases as a combination of tissue and tumor type, which we transformed to a compound string joined by a space (e.g., *Tissue: Breast* and *Type: Adenocarcinoma* became *Disease: breast adenocarcinoma*).

For **drugs**, we extracted the drug term from each knowledgebase and transformed it to lowercase text. As many interpretations contain more than one drug, we identified the delimiting character for each resource where multiple drugs are represented as a single string and split the string on the delimiter (e.g., the single string “*dabrafenib + trametinib*” was treated as the two strings “*dabrafenib*” and “*trametinib*”).

We did not perform this analysis for evidence levels, as there is no shared meaning behind unharmonized evidence levels across resources (**Table 1**).

### Project GENIE

GENIE data were downloaded from the 3.0.0 data release available online at: https://www.synapse.org/#!Synapse:syn7222066/files/. Variants were extracted from “data_mutations_extended.txt”, and clinical data from “data_clinical_sample.txt”. Variants were filtered on predicted consequence of medium or high impact. This classification was based upon the VEP consequence table (http://useast.ensembl.org/info/genome/variation/prediction/predicted_data.html#consequences) and resulted in exclusion of variants classified as *Silent*, *3’Flank*, *3’UTR*, *5’Flank*, *5’UTR*, *Intron*, or *Splice_Region*. Patients without any variants after filtering were included in all calculations. Oncotree xrefs were obtained from their API at http://oncotree.mskcc.org/api/tumorTypes (data version *oncotree_2018_05_01*), and xrefs were then mapped to DO terms where they matched. In cases where 1-to-many mappings occurred, manual review of those mappings was performed to select the most appropriate mapping.

### Variant intersection search

Variant coordinates were used to search genomic features via coordinate intersection. A complete intersection of query and target is considered a *positional match*, or a more specific *exact match* if the alternate alleles also match. A *focal match* is reported if the intersection fraction is less than complete, but over 10% overlapping (reciprocally). A *regional match* is reported if there is any intersection, but the match is of no other type (**Figure S4a**).

### Disease TopNode search

Disease searching returns a distance of the number of ancestor or descendent TopNode terms between the queried disease and the matching target. Two diseases sharing a TopNode term (e.g. *DOID:3008 - Invasive ductal carcinoma* and its parent term *DOID:3007 - Breast ductal carcinoma* both are members of *DOID:1612 - Breast cancer*) would have a distance of 0. However, if two diseases share a TopNode term but do not have a direct lineage, they are not a match (e.g. *DOID:0050938 - Breast lobular carcinoma* does not match to *DOID:3007 - Breast ductal carcinoma* even though they share a TopNode term (*DOID:1612 - Breast cancer*), as they are sibling concepts and do not have an ancestor/descendent relationship (**Figure S4b**).

### Gene intersection search

To assess cohort interpretability (**Figure S6**) when considering only matching a variant to a gene, we used the assigned gene symbols for each GENIE variant and compared them to interpretation gene symbols. Patients with at least one variant matching an interpretation gene symbol were considered a match. Matches were subsequently filtered by broad disease matching and by interpretation tier; no adjustment was made to the evidence level and tier to account for this imprecise aggregation strategy.

### ElasticSearch API and web frontend

Harvesters create *Association* documents segmented by the *source* field. Documents are posted to an ElasticSearch 6.0 instance provisioned by AWS elasticsearch service. Index snapshots are archived online: https://s3.console.aws.amazon.com/s3/buckets/g2p-ohsu-snapshots.

On top of Elasticsearch, we built web services using the Flask web framework. The search.cancervariants.org endpoint provides two simple REST-based web services: an association query service and a GA4GH beacon service. The association query service allows users to query for evidence using any combination of keywords, while the beacon service provisions G2P associations into the GA4GH beacon network (beacon-network.org) enabling retrieval of associations based on genomic location. OpenAPI (swagger) documentation is provided to accelerate development and provide API integration scaffolding. Client applications can use the API to create higher level sets of queries driven by cohort allele sets (e.g., MAF/VCF files) with varying genomic resolutions and disease/drug combinations. The API server and nginx proxy are described by Docker configurations and deployed co-located within a t2.micro instance.

The UI is a customized Kibana dashboard which enhances Lucene-based full-text search of associations with interactive aggregation heatmaps, tables and other components. The API documentation is available online at: search.cancervariants.org/api/v1/ui/

### Python interface and analysis notebook

The python 3.6 interface package and jupyter analysis notebook to generate these results are available online at http://git.io/vicckb.

### Data availability

Analyzed harmonized data from the aggregated knowledgebases are available for bulk download online at https://s3-us-west-2.amazonaws.com/g2p-0.10/index.html. Data are made available according to the data sharing principles and data sharing agreement provided by the VICC (online at: cancervariants.org/join). In accordance with these principles, all content is available for academic research. The CIViC, CGI Biomarkers, and PMKB knowledgebases provide content with no restrictions on reuse; however, commercial use of content from other knowledgebases is restricted—see individual knowledgebases for current content licensing. All code is open-source (MIT licensed) and available online at github.com/ohsu-comp-bio/g2p-aggregator (website) and git.io/vicckb (python interface).

